# Late-acting self-incompatible system, preferential allogamy and delayed selfing in the heteromorphic invasive populations of *Ludwigia grandiflora subsp. hexapetala*

**DOI:** 10.1101/2021.07.15.452457

**Authors:** Luis O. Portillo Lemus, Marilyne Harang, Michel Bozec, Jacques Haury, Solenn Stoeckel, Dominique Barloy

**Affiliations:** DECOD, (Ecosystem Dynamics and Sustainability), Institut Agro-Agrocampus Ouest, IFREMER, INRAE, Rennes, France; IGEPP, INRAE, Institut Agro, Univ Rennes, 35653, Le Rheu, France

**Keywords:** Mating system, Pollen tube elongation, Self-fertilization, Delayed selfing, Water primrose, Onagraceae, Myrtales

## Abstract

Breeding system influences local population genetic structure, effective size, offspring fitness and functional variation. Determining the respective importance of self- and cross-fertilization in hermaphroditic flowering plants is thus important to understand their ecology and evolution. The worldwide invasive species, *Ludwigia grandiflora subsp. hexapetala* (Lgh) presents two floral morphs: one self-compatible short-styled morph (S-morph) and one self-incompatible long-styled morph (L-morph). In this study, we identified the breeding systems of western European experimental and natural populations of Lgh by comparing structural characteristics of pollen and style, by studying self- and cross-pollen tube elongations and the viability of the resulting seeds and seedlings in both floral morphs. Our results showed no differences in pollen shape and stigma surfaces no matter the floral morph. In the self-incompatible L-morph flowers, self-pollen tubes were stopped tardily, in the ovarian area, and were unable to fertilize the ovules. This first formal identification of a late-acting, prezygotic self-incompatible system (LSI) in *Ludwigia* genus contributes a case of LSI in an additional family within the Myrtales order. In the self-compatible S-morph flowers, self-pollen always succeeded to self-fertilize the ovules that nearly all developed into viable seedlings. However, cross-pollen tubes always elongated faster than self-pollen tubes in both morphs. S-morph individuals may thus advantage preferential allogamy over selfing when cross-pollen is available despite its self-compatibility. As expected in late-acting self-incompatible systems, L-morph flowers authorised 0.2‰ of selfed seeds during the uppermost flowering season, that increased to 1‰ at the end of the flowering season. Such delayed selfing resulted in a significant quantity of viable floating seeds. They may contribute to the local regeneration, seed bank and propagation of the L-morph, which may contribute to explain its invasion success worldwide. Management plans of Lgh would gain to consider the breeding systems we identified.

## Introduction

Breeding systems are the main factor influencing the evolution of genetic diversity in populations and species with potential consequences for adaptation and speciation (Duminil et al. 2007; Ellegren and Galtier 2016). Around 50% of angiosperm species develop a variety of self-incompatible mechanisms that avoid self-fertilisation and favour outcrossing (Igic et al. 2008; Fujii et al. 2016). Self-incompatibility (SI) is the inability of functional male and female gametes to achieve self-fertilization. Prezygotic SI involve biochemical reactions resulting from the pollen-pistil interaction that blocks incompatible pollen before sperm fertilizes an egg. Out of a handful of model species, the precise nature of these reactions in most species is not clearly understood, with an intriguing paradox: recurrent patterns of SI are ubiquitous across families but with a large variation in the type of sites of reaction (Charlesworth et al. 2005; Allen and Hiscock 2008; Shimizu and Tsuchimatsu 2015). Identifying the site of the incompatible pollen-pistil rejection is an essential step to understand and predict how its breeding system may shape the evolution of its populations in their ecological contexts (Charlesworth et al. 2005; Takayama and Isogai 2005; Busch and Schoen 2008; Ferrer and Good 2012; Grossenbacher et al. 2017). Moreover, characterizing the prevalence of SI systems developed in species, genera and families across the Angiosperms and eukaryotes contribute understanding which biological, ecological and evolutionary features may explain the ubiquitous occurrence of mechanisms favouring allogamy across the tree of life (Charlesworth et al. 2005; Igic et al. 2008; Santos-Gally et al. 2013; Gibbs 2014; Fujii et al. 2016; Grossenbacher et al. 2017; Barrett 2019).

We currently consider three major mechanisms of SI in flowering plants, homomorphic gametophytic, homomorphic sporophytic and heteromorphic sporophytic, that differ in the site of the rejection of incompatible pollen tubes (Gibbs and Bryan 1986; Dickinson et al. 1992; Hinata et al. 1993; Barrett and Cruzan 1994; Kao and McCubbin 1996; de Nettancourt 1997). Homomorphic SI systems imply that flowers with different SI types present the same flower shape and structure. By contrast, heteromorphic self-incompatibility (HetSI) associates different compatibility between individuals with different floral morphologies, sometimes associated with additional features such as different patterns of spatial separation of anthers and stigmas (Webb and Lloyd 1986; Opedal 2018), differences in pollen sizes and shapes, and different lengths of stigmatic papillae (Darwin 1877; Barrett and Shore 2008; Igic et al. 2008; Barranco et al. 2019; Barrett 2019; Matsui and Yasui 2020). Commonly, species with style polymorphism have a sporophytic heteromorphic (*i.e*., di- or tri-allelic) incompatibility system that prevents self-fertilization and crosses between individuals of the same floral morph (Barrett 2019).

In most studied species developing these three SI systems, pollen-tube growth is stopped from the stigma or in the style as a result of a rapid response to the pollen–pistil interaction. However, some species present a delayed reaction of self-pollen rejection, named ovarian or late-acting self-incompatibility (LSI) because it occurs lately in the ovarian area (Seavey and Bawa 1986; Gibbs and Bianchi 1993; Sage et al. 1994; Gibbs 2014). It often coincides with a residual permeability of their SI systems resulting in a steady low level of selfing despite an effective late-acting SI (Seavey and Bawa 1986; Gibbs 2014). It concerns both homomorphic and heteromorphic species (Gibbs 2014; Simon-Porcar et al. 2015). For example, the homomorphic LSI Theaceae species *Camelia oleifera* and *Camelia sinensis*, allow 10% and ~2% of self-fertilization resulting in viable seeds, respectively (Chen et al. 2012; Liao et al. 2014). In the ovarian LSI heteromorphic *Narcissus spp*. from the Amarilidacea, from 4 to 30% of their seed-sets result from self-fertilization (Barrett et al. 2004; Medrano et al. 2012; Simon-Porcar et al. 2015). Within pre-zygotic LSI, Gibbs (2014) proposes to distinguish LSI mechanisms stopping pollen tubes before ovaries (as found in *Melaleuca spp*. and *Thryptomene calycina*, Barlow and Forrester 1984; Beardsell et al. 1993) from those stopping self-pollen tubes while penetrating the ovules (as found in *Acacia retinodes*, Kenrick et al. 1986).

In the *Ludwigia* genus (Onagraceae), 8 of 82 species have been reported as “self-incompatible”, although the nature of their SI were not formally studied and established (Raven 1979). The water primrose, *Ludwigia grandiflora* subsp. *hexapetala* (Hook. & Arn.) Nesom and Kartesz (2000), (hereafter *Lgh*), is one of the most invasive aquatic plants in the world, currently spreading out of South America into North America, Europe and Eastern Asia (EPPO 2011, Thouvenot et al. 2013; Portillo-Lemus et al. 2021a). Tackling how this species reproduces is crucial for understanding its expansion and alleviate its impacts in aquatic environments (EPPO 2011). Yet, we still don’t know how sexual reproduction works in this invasive species (Dandelot 2004; Ruaux 2008; Thouvenot et al. 2013). *Lgh* presents two types of floral morphologies in European populations: a short-styled morph (S-morph) that is self- and intra-morph compatible (crosses produce viable seeds), and a long-styled morph (L-morph) that is self- and intra-morph incompatible (crosses do not produce seeds, Portillo-Lemus et al. 2021a). In both floral morphs, inter-morph crosses always produce viable seeds. The two floral morphs of *Lgh* develop bowl-shaped non-tubular flowers with two whorls of stamens of different heights. These two characteristics contrast with typical heteromorphic SI species that most frequently present tubular flowers and only one whorl of stamens (Barrett and Shore 2008; Cohen 2010; Barrett 2019). Interestingly, floral morphs are mostly found in allopatric monomorphic populations (*i.e*., exclusively S-morph or exclusively L-morph populations) in Western Europe and other invasive worldwide populations (Hieda et al. 2020; Portillo-Lemus et al. 2021a). Surprisingly, around 75% of the invasive populations worldwide seem exclusively composed of self-incompatible L-morph individuals, tackling the paradigm that successful invasive species should be the ones capable of self-fertilization (Baker 1955; Cheptou 2012; Razanajatovo et al. 2016). Recently, some isolated monomorphic L-morph self-incompatible populations were however found to produce seeds (Portillo-Lemus et al. 2021a). These puzzling observations question the type of SI reaction involved in the self-incompatible L-morph and its level of permeability to self -fertilization. More broadly, they question the type of breeding systems developed in these invasive populations (Hieda et al. 2020; Portillo-Lemus et al. 2021a).

In this study, we thus aimed at better characterizing the breeding system of the two floral morphs in Western European populations of *Lgh* by tracking the fates of self- and inter-morph pollen tubes in both floral morphs. First, we searched for structural evidences through analysis of the morphologies of pollen grains and stigma surfaces of the two floral morphs that may be characteristic of homomorphic and heteromorphic sporophytic self-incompatibilities. Second, to better categorize the type of the SI developed by this species, we aimed at identifying the site where the incompatible pollen tubes were blocked in the self-incompatible L-morph pistils. To achieve this goal, we followed the germination and the progression of the self- and inter-morph-pollen tubes on histological sections, prepared at different times after pollination. Third, to quantify the permeability of this self-incompatible system, we counted the number of viable seeds and seedlings obtained from self-fertilization in controlled conditions and in natural monomorphic L-morph populations. Fourth, to test if the S-morph individuals truly reproduced using selfing, as expected in this self-compatible morph, we followed the progression of the self- and inter-morph-pollen tubes, as described for the morph-L, and studied the viability of obtained seeds and seedlings. Finally, we discuss on the contribution of the patterns of SI reactions and breeding system found in *Lgh* with those already observed in other species, with a special focus on the Myrtales order that includes the *Ludwigia* genus.

## Material and Methods

### Plant material and sampled populations

To better characterize the breeding system of the two floral morphs that were previously reported in *Lgh*, (Portillo-Lemus et al. 2021a), we studied a total of 100 individuals growing along the Loire valley in France (Table S1 for GPS location). We sampled five monomorphic populations only composed of S-morph individuals (distance between them: min=40km, mean=146km, max=301km) and five monomorphic populations only composed of L-morph individuals (distance between them: min=6km, mean=124km, max=274km). In each population, we randomly sampled 10 stems per population. *Lgh* partly reproduce using clonal budding and rhizomes (Thouvenot et al 2013). We also know using allele sharing distances between pairs of individuals from 30 individuals genotyped using 38 polymorphic SNPs that, at least in one of these monomorphic populations (Mazerolle), populations present different genotypes within a same floral morph though with a restricted number of ancestors (Genitoni et al. 2020). Sampled stems of all S-morph and L-morph populations were transplanted into a common greenhouse (location: L’institut Agro, Rennes, France. 48°06’47.7”N 1°42’30.2”W) where individuals from one population were growth together in an 80L container to avoid environmental influences on pollen shapes, pollen tube elongations, ovule and seed developments. The 10 containers were watered every 15 days with a commercial nutrient solution (6% nitrogen, 6% phosphorus and 6% potassium) during growth and flowering periods to avoid nutritional deficiencies.

### Measure of herkogamy

To link breeding systems with floral morphs, we measured the spatial separation on a vertical axis of anthers and stigmas within flowers (herkogamy) from the 10 *Lgh* populations we sampled, as this floral trait would have a negative relationship with the rate of autofertility in plant species (Webb and Lloyd 1986). Herkogamy is quantified as the difference between pistil and stamen lengths (respectively, 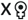 and 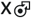) with their natural twists and curvatures (Opedal 2018). Flowers with stigmas positioned above or protruding beyond the anthers would have higher probability to be contacted first by a visiting pollinator which corresponds to approach herkogamy. Flowers in which anthers are above the pistil and with higher probability to be contacted first by pollinators which corresponds to reverse herkogamy. *Lgh* flowers present two whorls of stamens. We thus did 150 measures, i.e., the length of the pistil 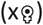 and the length of the inner 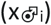 and outer 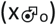 stamens per floral morph, and calculate difference between length of pistil and length of anther.

### Histological preparations

To observe pollen and stigmatic papillae morphology, to follow pollen tube growth and to achieve histological sections of ovaries for studying embryo development, we fixed flowers in FAA 1: 2: 20 (formalin: acetic acid: ethanol V/V). Pollen germination, pollen tube growth and ovule penetration were observed using the aniline blue fluorescence method (Martin 1959). The fixed samples were washed several times with distilled water and each flower was placed in 1mL of staining solution (0.1% w/v aniline blue, 1% v/v tween-20^(R)^, 0.2 M K_2_PO^4^, 0.1 M NaOH) for 1 hour at 95°C. After squashing the pistils between the slide and coverslip, observations were made under ultraviolet light (335-364 nm) with a microscope Leica DM4000 B©, and a camera 190HD. To observe embryo development, fixed ovaries were dehydrated, embedded in paraffin, sectioned at 15 μm with a microtome, and mounted on glass slides (Sakai, 1973). Sections were placed in 0.05% toluidine blue O in distilled water for 2–30 min, rinsed once in water for 1 min, and air-dried. Paraffin was removed using two xylene baths and the cover slip was mounted with resin. Histological preparations were realized by Nublat Laboratory (https://www.laboratoire-nublat.com/118botanique.html).

### Pollen and stigmatic papillae morphologies

To assess if the two floral morphs found in western Europe populations of *Lgh* showed difference in pollen shapes and sizes in association with floral morphs, we studied the shape of 500 pollen grains per floral morph. For each morph, we sampled 25 flowers in the 5 corresponding populations. For each of these flowers, we measured 20 pollen grains from a mix of the stamen verticels, resulting into a total of 1000 pollen grain measures for both morphs. We measured their diameter under an optical microscope. To assess if S-morph and L-morph flowers of *Lgh* showed difference in stigmatic papillae structure as sometimes found in some heteromorphic SI species, we also noted the long or short shape of stigmatic papillae in 30 histological stylar sections for each of the two floral morphs (Dulberger 1992).

### Pollen tube growth, ovule fertilisation and embryo-sac development

To identify in the self-incompatible L-morph pistil where self-pollen tubes were blocked and to characterize the type of breeding preferentially achieved in the two floral morphs, we performed hand-controlled pollinations to track and compare the self- and inter-morph pollen tube growth in both floral morphs. As experimental pollinations in *Lgh* have previously shown that self- and intra-morph pollinations in L-morph flowers gave similar rates of rejections (Portillo Lemus et al 2021), here we focused our experiments on a comparison between self-pollinations (pollen grains from the same flowers) and reciprocal inter-morph pollinations. To perform controlled self-pollinations, we enclosed early flower buds in cellophane bags to protect them from external pollen. To effect self-pollination, we dissected mature anthers using tweezers to place pollen directly on its own receptive stigma and re-enclosed the flowers in cellophane bags. For reciprocal inter-morph pollination, to simulate free random crosses, we selected five pollen-donor flowers from the other morph to generate a pollen mix. Flowers to receive inter-morph pollen were emasculated before anthesis and then pollinated with this mix of pollen. After pollination, the pollinated flowers were once again enclosed in cellophane bags in order to protect them from any contaminant pollen.

To study self- and inter-morph pollen tube growth in both floral morphs, we made 150 self-pollinations per morph and 300 reciprocal inter-morph pollinations between S-morph and L-morph. At 2, 3, 7, 16, 24 hours after the hand-controlled pollination, we randomly sampled and fixed 30 flowers per morph and pollination type. For each flower, pollen tube growth and ovule fertilisation were observed using aniline blue fluorescence method with squashed pistils. We measured the growth of the pollen tubes, *i.e*., the distance between the stigma and the tip of pollen tube (d, μm) at 2, 3, 7, 16, 24 hours (*t*) after the hand-controlled pollinations. From these measures, we calculated the speed of pollen tube (speed) between two measures in micrometres per hour (μm/h) as 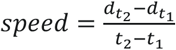 and the acceleration of pollen tube growth in micrometres per hour square (μm/h^2^) as 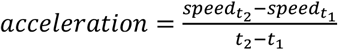.

To evaluate the viability of fertilisation and study the embryo-sac development, we analysed a total of 2520 ovary sections corresponding to 14 sections of five flowers per floral morph and cross type, over nine-time steps (1, 2, 3, 4, 5, 6, 7, 15, 21 days after pollination (*dap*).

### Fruit and seed set production

To assess the incidence of self-fertilisation in the self-incompatible L-morph individuals, 1600 self-pollinations were carried out from July to August (summer), and from September to October (fall), respectively. Self-pollinated L-morph flowers were randomly selected within groups of individuals growing in the greenhouse. As a control, we made 30 self-pollinations with self-compatible S-morph flowers, 30 L x S and 30 S x L-morph pollinations during summer and repeated this sequence during the fall.

We also assessed the rate of self-fertilisation in self-incompatible L-morph from *in situ* populations at the beginning of October. We collected the fruits of five S-morph and five L-morph *in natura* populations to evaluate the fruit and seed sets. To quantify the fruit-set *in natura*, we counted fruits produced in five quadrats of 1 m^2^, 10 m away on a linear transect. As S-morph populations showed a massive fruit production, we estimated the seed-set production per m^2^ as the mean number of seeds produced by 25 fruits randomly picked in the quadrats (5 fruits per quadrat). As the L-morph *in natura* populations produced few fruits, we collected and counted all the seeds produced in all the fruits found in quadrats. We sowed all the *in-natura* produced seeds in the greenhouse to calculate the germination rate as the number of seedlings over the number of sowed seeds.

### Statistical analysis

To examine possible differences in pollen tube elongations at each time step (time=[2, 3, 7, 16, 24 hours]), and the changes in speed and acceleration of elongations along time in the styles of the two floral morphs considering the four possible types of crosses (cross_types=[S-morph self-pollination; S-morph x L-morph; L-morph self-pollination; L-morph x S-morph]), we used ANOVA (type = III) tests. For each time step, we computed three ANOVA models: pollen_tube_length~cross_types, speed~cross_types*time and acceleration~cross_types*time. For each test, we verified the homoscedasticity of distributions between groups and the normality of residuals using Shapiro–Wilk’s tests. When ANOVAs were significant, we applied Tukey’s honestly significant difference (HSD) posthoc comparisons of the means to identify which groups and parameters caused those significant differences. We compared measures of herkogamy between S-morph and L-morph flowers using a Mann-Whitney U test and discussed considering their compatibility and realised breeding systems. All analyses were performed using the stats package in R 4.0.4 software (R Development Core Team, 2014). All results were summarised in the supplementary information table S2.

## Results

### Measures of herkogamy

The means and standard deviations of measures of herkogamy (pistil length – stamen length in a same flower, Figure S1) for L-morph flowers outer whorls were of +2.00±0.65mm and +0.72±0.35mm for the inner whorls. In S-morph flowers, they were of +0.85±0.64mm for the outer whorls and of −1.50±0.43mm for the inner whorls. In the L-morph flowers, the heights of the anthers of the 150 outer and 150 inner measured whorls were, respectively always (U=22500, p<0.001) and significantly (U=21050, p<0.001) below the heights of the pistil. In the S-morph flowers, the heights of the anthers of the 150 outer measured whorls were significantly below the heights of the pistil (U=19930, p<0.001), while the heights of the anthers of the 150 inner measured whorls were all above the heights of the pistil (U=22500, p<0.001).

### Pollen and stigmatic papillae morphologies

Pollen grains were all released in triporate monads and connected to the anthers by viscin threads (Fig.1; a3). In both floral morphs and in all the studied populations, pollen grains were all smooth and had the same sizes ranging from 60 to 95 μm whatever their floral morphs (ANOVA, p-value=0.228, Fig. 1 b3, c3). The styles of both floral morphs had capitate stigmas, which were all wet during anthesis and had stigmatic papillae which were submerged by mucilage (Fig. 1; a1, b1). In both floral morphs, all the 60 longitudinal sections of the stigma receptive surface (30 per morph) showed the same morphology of elongated unicellular papillae (Fig. 1; a2, b2). The only difference we found between styles of floral morphs concerned their lengths: styles of S-morph flowers were significantly wider (2.27mm ± 0.19) and shorter (8.04 mm ± 0.19) than those of L-morph flowers (width: 1.53 mm±0.09; length: M=8.97±0.25; p-value<0.001, Fig.1; c1, c2), resulting in a difference of around 1mm between long- and short-style lengths.

**Figure 1:**
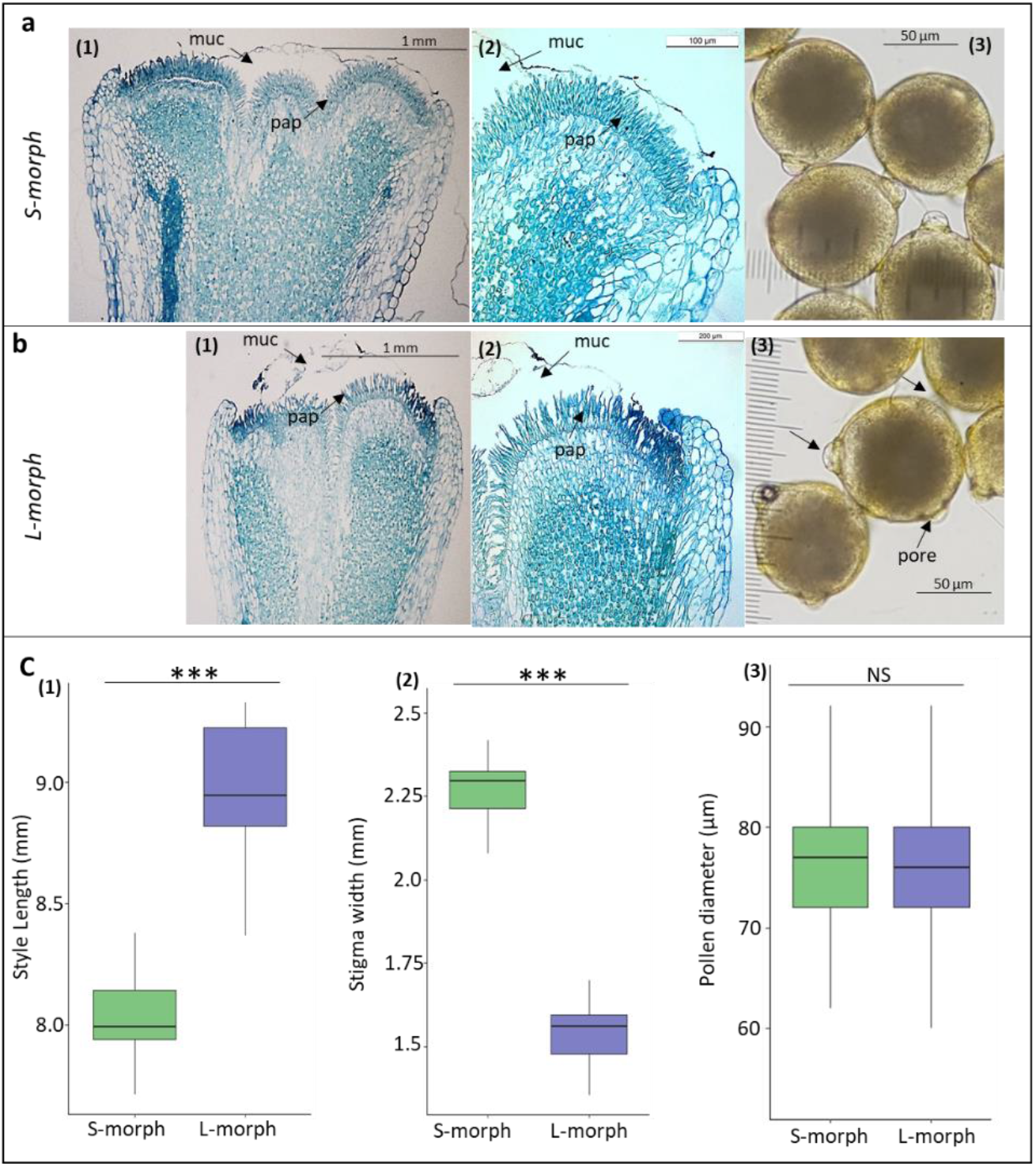
Stigma and pollen morphologies in both floral morphs of *Ludwigia grandiflora subsp. hexapetala*. Stigma section of S-morph (a 1, 2) and L-morph (b 1, 2) under the 10x (a1, b1) and 20x (a2, b1) objectives of optic microscope stained with toluidine blue O. Stigma sections of the stigma have a thickness of 15 μm. In both floral morphs, the stigma was filled with mucilage (mu) (a1-2, b1-2,). Both floral morphs showed the same shape of stigmatic papillae (pap). - Pollen morphologies of S-morph (a3) and L-morph (b3) under 20X objective of optical microscope. Both morphotypes shared the same shape and size of pollen (a3, b3) and the same diameter of pollen (c3).

### Pollen tube growth and fertilisation

All pollen grains germinated, elongated down the style (Fig. 2a-c), and reached the ovules 24 hours after pollen deposition on stigmas (Fig. 2d-e), in self and inter-morph crosses and independent of the pollen origin. In the self-pollinated L-morph flowers, only six ovules of 300 studied (2 %) were penetrated by a pollen tube with pollen tube tips located in the area of the embryo sac (Fig. 2f). Figure 2f shows an ovule surrounded by several pollen tubes, but it is unclear whether any of them have penetrated the micropyle. The other 294 unpenetrated ovules showed tips of pollen tubes with a small bulge, stopped between the ovarian tissues and the beginning of the first cell layers of the ovule integuments and we found no traces of pollen tubes in the area of the embryo sac (Fig. 2d, e). All the 300 L-morph ovules that were cross-pollinated with S-morph pollen were penetrated by pollen tubes 24 hours after pollination. All S-morph ovules were penetrated by one pollen tube 24 hours after self and inter-morph pollinations (Fig. 2g).

**Figure 2:**
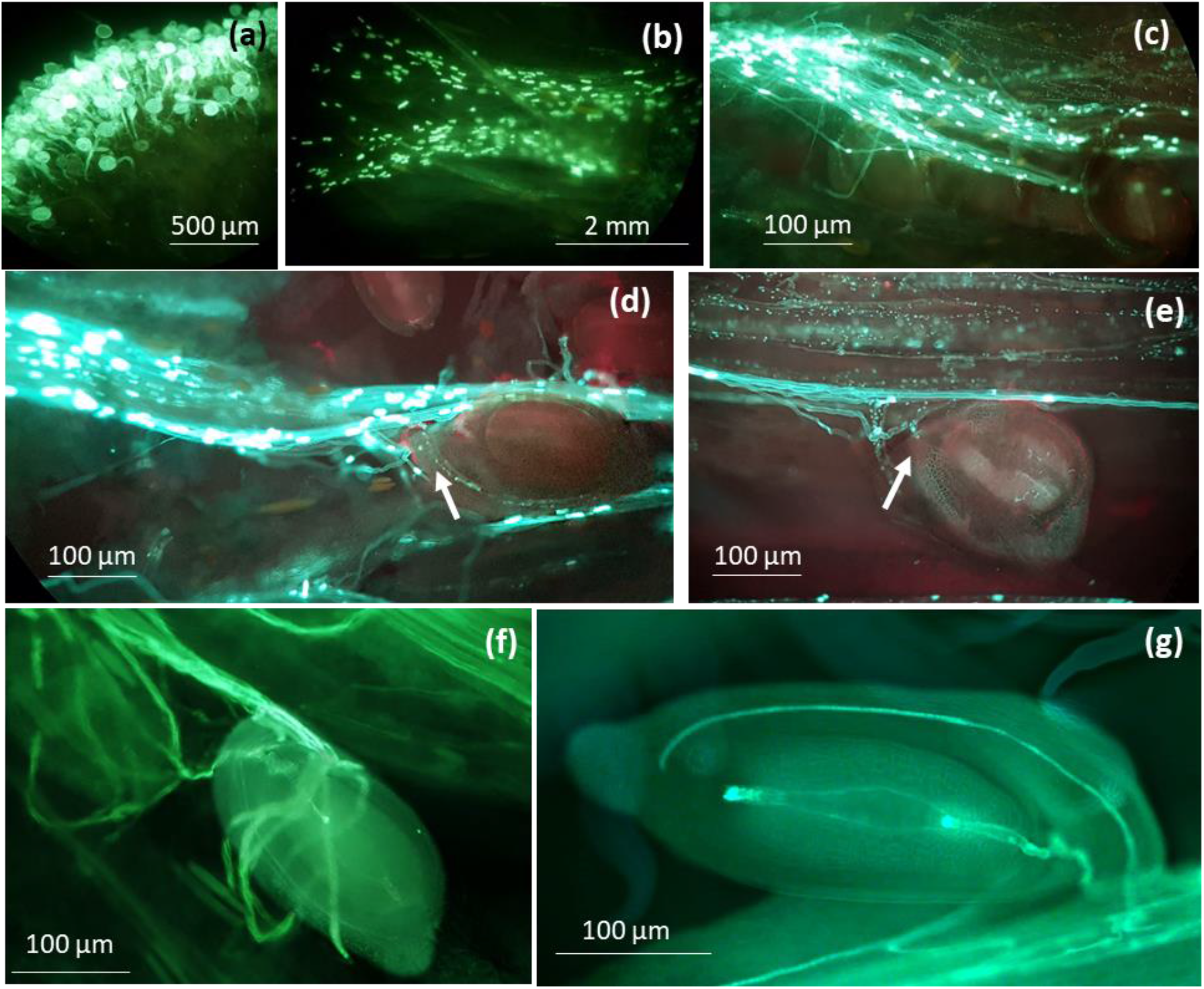
Fluorescence aniline blue staining of self-pollen tube in L-morph pistils (a-f). a) Pollen grains germination on a stigma surface (zoom 10X); b) Pollen tubes elongating in a style (zoom 2.5X); c, d, e) Pollen tubes in the ovarian area (zoom 20X); f) Pollen tubes in the area of embryo sac (noticed for 6 pollen tubes over 300 studied). g) Pollen tubes in the area of embryo sac in S-morph ovary after self-pollination.

### Kinetic elongation of pollen tube

In the styles of both floral morphs, until reaching the ovules, inter-morph pollen tubes outdistanced self-pollen tubes (HDS-test, p-value<0.001; Mean and SD values reported in Table S2; Fig. 3). Significant advances of inter-morph pollen tubes were observed from two hours after pollinations. (Fig. S2a). Speed of elongations of pollen tubes in both morphs quickly increased in the first two hours after germination while penetrating the stigma tissues to reach an elongation of 0.3-0.4 mm/hours all along the style, and then doubled their speed of elongation when entering the ovary area (Fig. S2b, Table S2).

**Figure 3:**
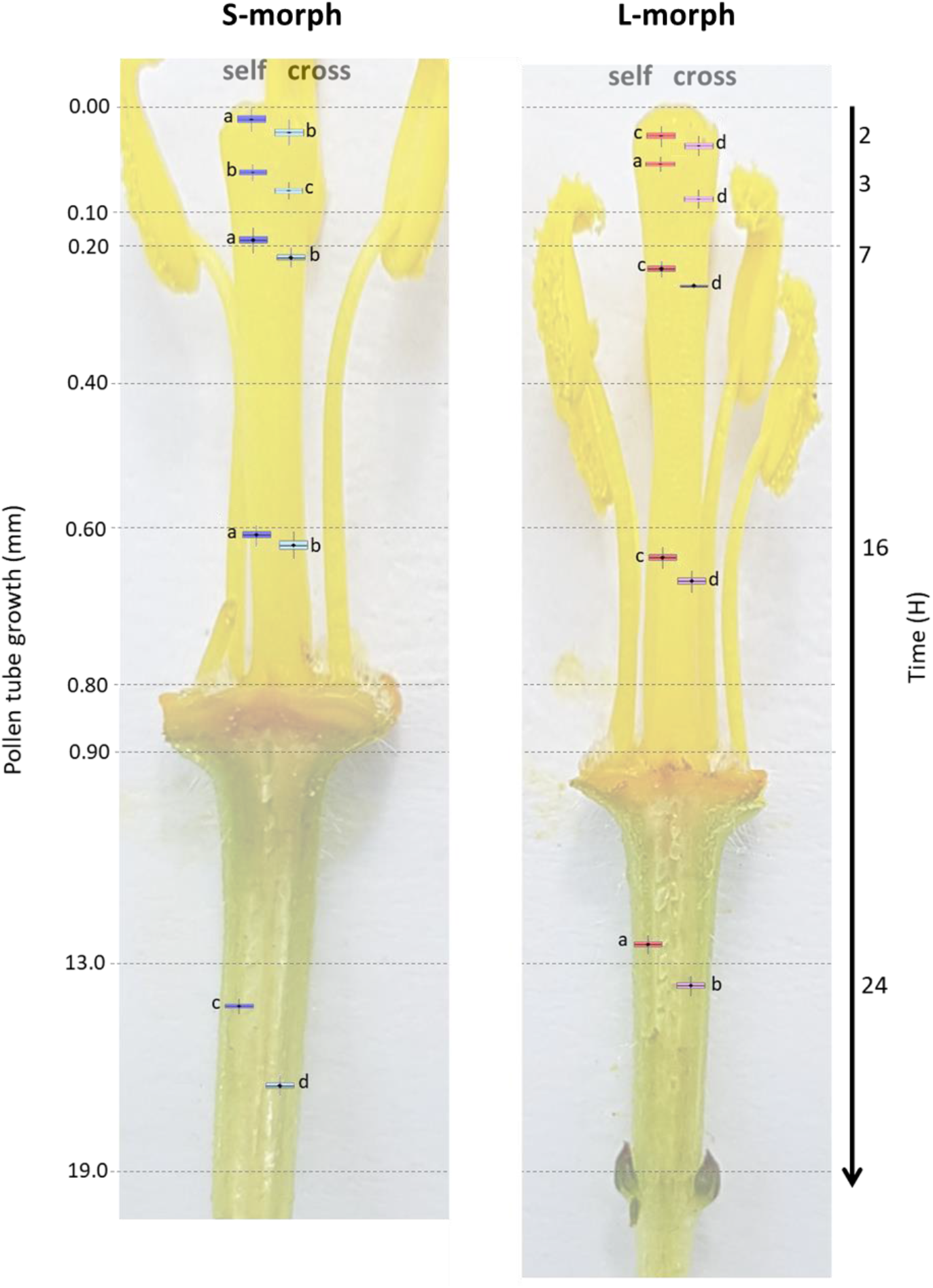
Kinetic of pollen tube elongations in both floral morphs in *Ludwigia grandiflora subsp. hexapetala*. a) Kinetic of pollen tube elongations in S-morph styles when self- and cross-pollinated (left, blue and sky-blue boxes respectively); b) Kinetic of pollen tube elongations in L-morph style when self- and cross-pollinated (right, red and pink boxes respectively). Significance letters were obtained using a Tukey’s honestly significant difference (HSD) post-hoc comparisons of the means. In both floral morphs, cross-pollen tubes elongated faster than self-pollen.

### Fertilisation and embryo development

In both floral morphs, each locule of the pentacarpellate ovary has 12 ± 2 ovules (Fig. S3). The anatropous and bitegmic ovules were disposed along a single longitudinal axis on placentae (Figs. 4, 5 and S3). The embryo sac (female gametophyte) was an *Oenothera-type* and composed of only 4 cells: 2 synergids and an egg-cell on the micropyle side, and a single polar nucleus in the central cell, matching previous observations made on other *Ludwigia* species and other Onagraceae genera (Fig. 4, 5; Rigakishi 1918; Eyde 1977; Tilquin and de Brouwer 1982; Tobe and Raven 1986).

**Figure 4:**
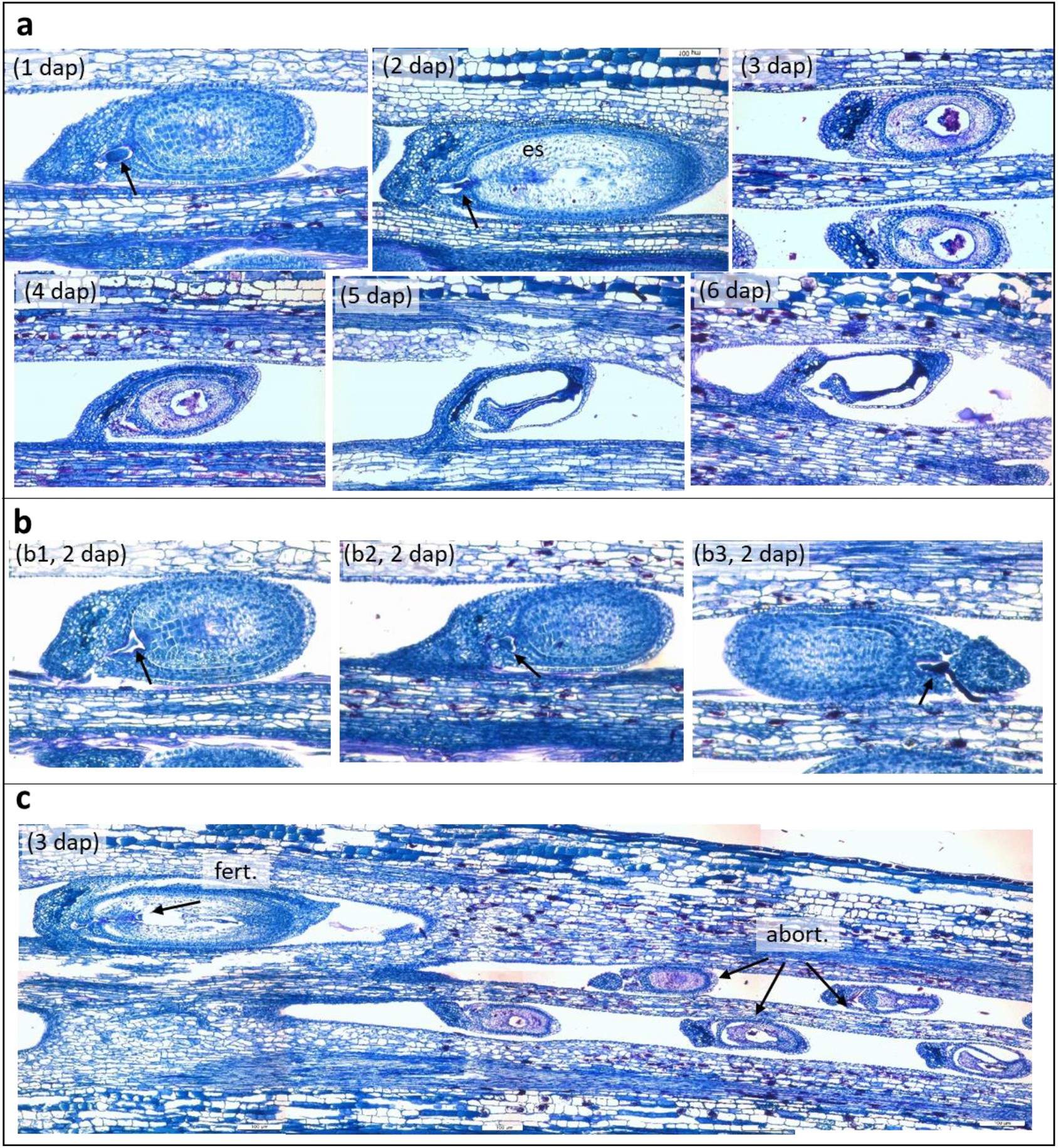
Temporal evolution of embryo sacs, ovules and ovaries in self-incompatible L-morph flowers after self-pollination. a) Abortive ovules sections after 1, 2, 3, 5 and 6 days after self-pollination (dap); b) Self-pollen tubes that stopped their elongations two days after self-pollination (dap) in three different ovules (1 to 3). The arrows indicate the tips of the self-pollen tube stopped between the ovarian tissues and the beginning of the first cell layers of the ovule integuments (see figure 5a for a comprehensive representation of the ovule parts); c) Example of a rare self-fertilisation event (fert.) obtained three days after self-pollination (dap) informing the “small fruit” formation. Only ~0.33% ovules were fertilised (1/300 ovules observations) (fertilisation = fert; abortive ovules= abort). All sections stained with toluidine blue O (zoom 10X). See figure 5 a comprehensive representation of the ovule parts.

**Figure 5:**
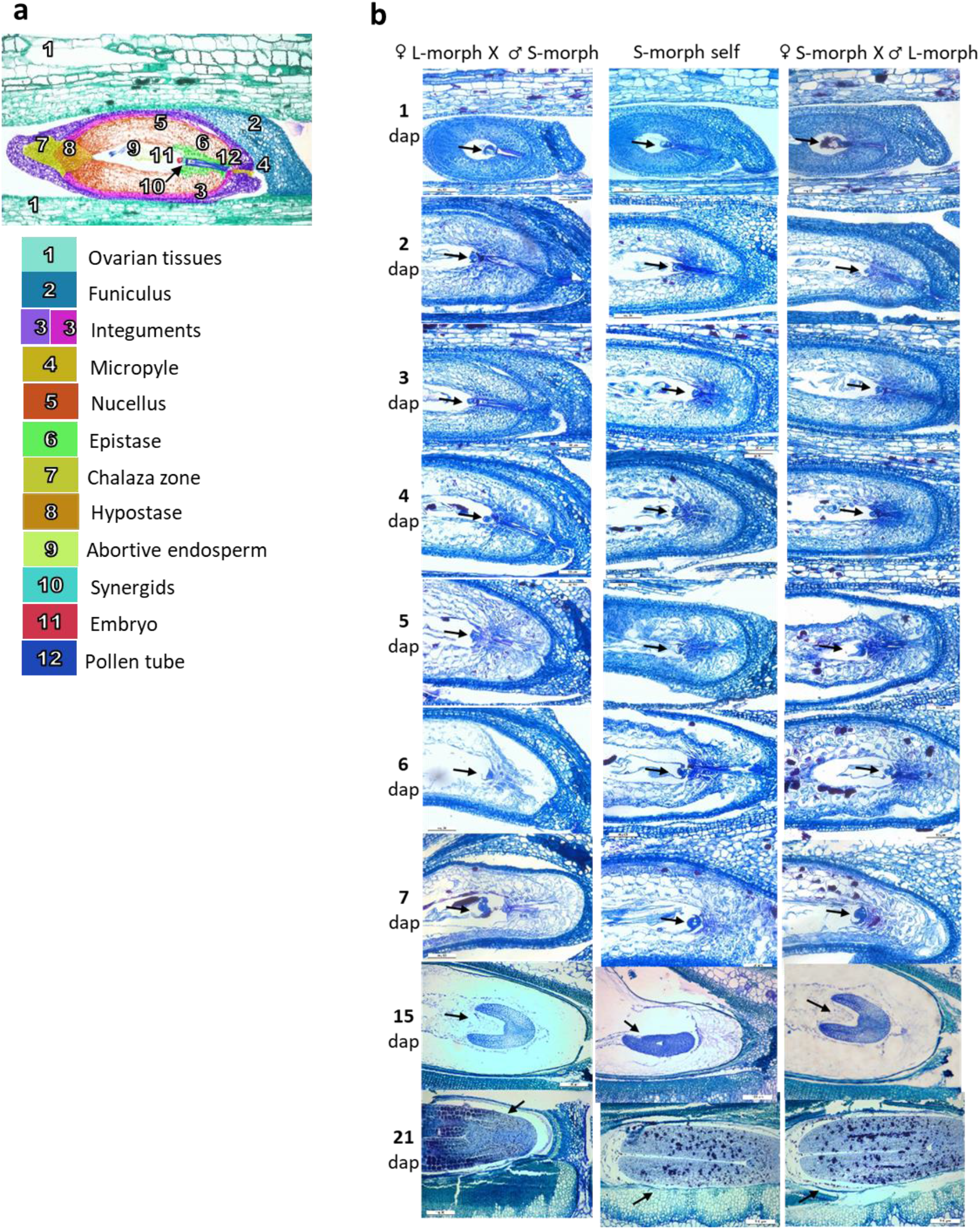
Fertilisation and embryonic development after cross and self-pollinating the self-incompatible L-morph and the self-compatible S-morph flowers. a) A comprehensive, colorized scheme of the different tissue parts of a L-morph ovule 3 days after disassortative pollination. b) Evolution of the ovules 1, 2, 3, 4, 5, 6, 7, 15, and 21 days after pollen deposition on the stigmas (dap): (1) L-morph ovules after cross-pollination 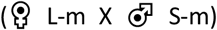; (2) S-morph ovules after self-pollination; (3) S-morph ovules after cross-pollination 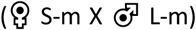. The arrows indicate the development of embryo. All sections were stained with toluidine blue O, 20X zoom under optical microscope.

In L-morph flowers, 24 hours after pollination, 98% of self-pollen tubes stopped their elongations, lately, in between the ovarian area and the beginning of the first integuments (Fig. 4.a, b). In these figures, the pollen tubes seem to be located before the nucellus-epistase out of the embryo sac and did not succeed to pass the synergids. However, it is always difficult to obtain sections showing all together, the synergids, egg cell and central cell which compromise the possibility to identify precisely where the pollen tube stopped. The embryo sacs found with self-pollen tubes stopped didn’t develop and degenerated three days after pollination (Fig. 4.c). In the greenhouse, self-pollinated L-morph flowers abscised three days after pollination which is consistent with the temporality of our histological observations (Fig. 4a). In comparison, the abscission of the unpollinated flowers of L-morphs occurred four days after emasculation, when protected with a cellophane bag. Under aniline blue fluorescence, we observed that 2% of self-pollen tubes (6/300) in L-morph flowers succeeded to reach the embryo sac and that only one self-fecundation event was observed over the 300 self-pollinated L-morph studied ovules from the ovary sections (Fig. 4b).

In S-morph, regardless of pollen origin, and in L-morph when inter-morph cross-pollinated, the first cell divisions of the embryos began two to three days after pollination (Fig. 5). From five days after pollination, the nucellus gradually began to disintegrate while the embryo continued to develop and grow. At 15 days, the nucellus disappeared completely, giving way to exalbuminous embryo development, which concluded with a viable seed 45 days after pollination.

### Fruit production in self-incompatible L-morph

In the greenhouse, during summer, only four very “small fruits” containing one, four, five and nine seeds respectively developed over 1600 studied L-morph flowers that were hand-pollinated with self-pollen only (Fig. 6). Considering that 60 ovules on average were initially available per flower, the rate of self-fertilisation was 0.02% (i.e., 19 seeds from 1600 x 60=96000 potential ovules available). In the greenhouse, during the fall, at the end of the flowering season, 25 very “small fruits” containing between one and ten seeds for a total of 98 seeds (Fig. 6), gave a rate of self-fertilisation of 0.1% (98 on 96000 potential ovules), thus 5-fold more than obtained in July, during the high fruiting period. All 30 S-morph flowers, which were self-pollinated as a control, developed into 30 fruits containing a total of 1800 seeds, with a constant number of 60 seeds per fruit both in July and September (Fig. 6).

**Figure 6:**
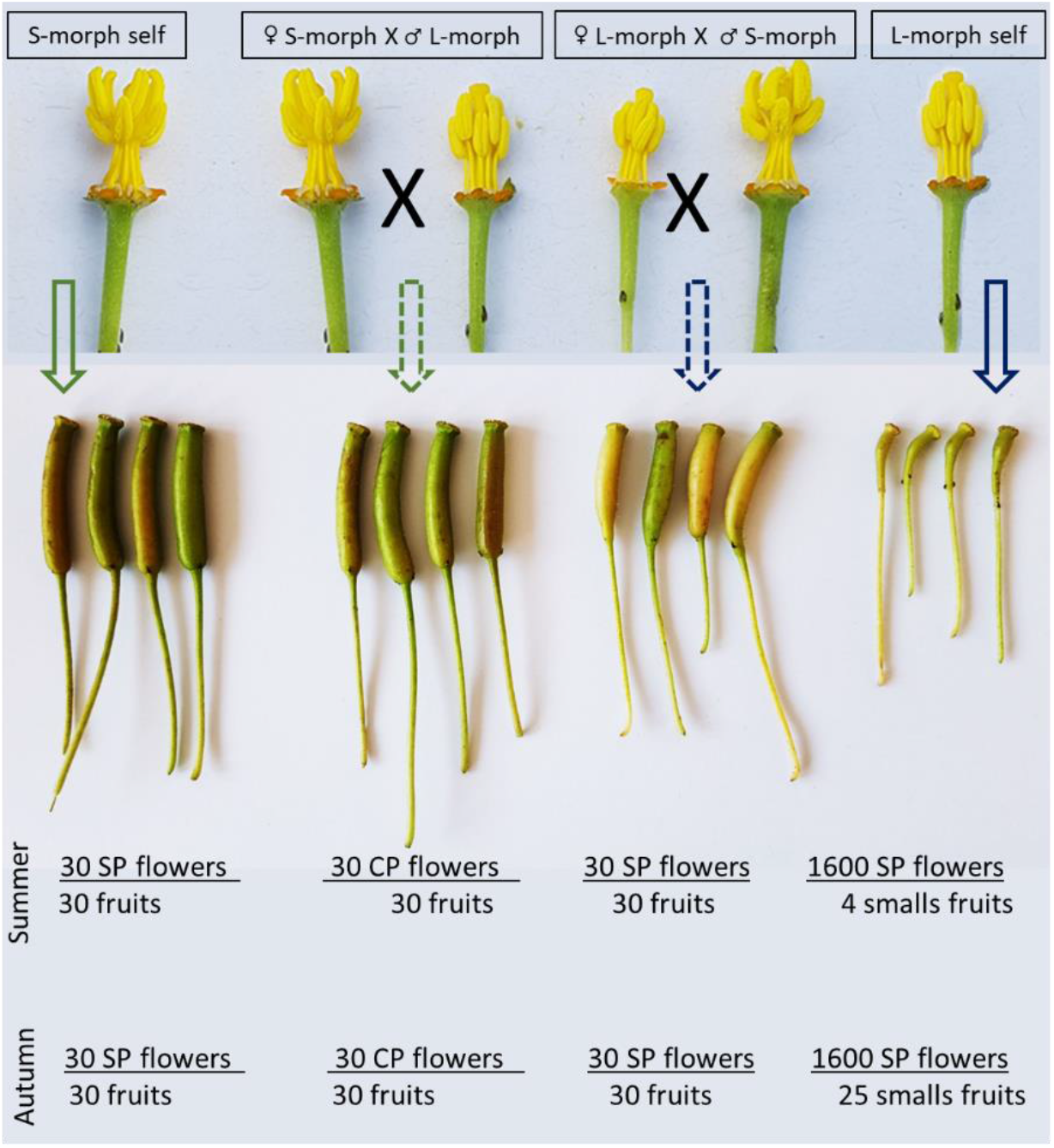
Summary of the fruit-set obtained from S-morph and L-morph flowers after hand-controlled pollination (SP = Self-pollination; CP = Cross-pollination) in summer and autumn. All S-morph flowers produced fully-seeded fruits, in summer and autumn, both after self- and cross-pollinations. All L-morph flowers produced fully-seeded fruits, in summer and autumn, when pollinated with S-morph pollen (L-morph x S-morph). 0.25% and 1.56% of L-morph flowers produced “small fruits” in summer and autumn when self-pollinated. These “small fruits” contained one to nine seeds that all successfully germinated and grew into viable seedlings.

To validate the results obtained in experimental conditions, we then assessed the production of “small fruits” in five L-morph and five S-morph *in natura* populations (Table S1). We found between 15 and 35 “small fruits” per m^2^ in self-incompatible monomorphic L-morph populations. All the “small fruits” collected *in natura* contained one to 15 seeds per fruit, as we obtained in experimental conditions in greenhouse. We counted a production of 92 to 217 seeds per m^2^. In the five self-compatible S-morph populations, we counted 384 to 864 fruits per m^2^. On five of these fruits per population, we counted 50 to 70 seeds per fruit, giving an estimation of 19200 to 51840 seeds per m^2^ (Table S1). All seeds from “small fruits” obtained in greenhouse and *in natura* germinated, and developed into viable seedlings (Table S1).

## Discussion

The results we obtained from a common garden and natural populations showed that both floral morphs found in European invasive populations of *Lgh* reproduced using a mixed breeding system, with mechanisms that may result in the production of allogamous and delayed autogamous viable seeds. Pollen grains and stigma surfaces of both floral morphs showed similar sizes and shapes. In the self-incompatible L-morph flowers with approach herkogamy, self-pollen tubes were only stopped when reaching the ovaries, before penetrating the ovules. These observations argue for a prezygotic ovarian LSI system in L-morph flowers of *Lgh*, corresponding to Gibbs (2014) classification. As commonly observed in this kind of SI system (Seavey and Bawa 1986; Gibbs 2014), a small proportion of self-pollen (0.02%), increasing by a factor of 5 at the end of the flowering seasons, succeeded to fertilize the ovules and to develop into seeds. All these seeds then developed into viable seedlings. In the self-compatible S-morph with reverse herkogamy, self-pollen grains succeeded in fertilizing the embryo-sacs that successfully developed into viable seeds and then into viable seedlings. Yet, in the two *Lgh* floral morphs, inter-morph pollen tubes always elongated faster than self-pollen tubes, which may give advantage to intermorph crosses when intermorph pollen is available.

### The prezygotic LSI system in Lgh: a rare but already observed SI system

European invasive populations of *Lgh* displayed two floral morphologies, a long-styled morph (approach herkogamy) with a prezygotic LSI and a short-styled morph (reverse herkogamy) that was self-compatible, with similar pollen shape and papillae structure regardless of the floral morph.

Previous literature may lead to associate HetSI with differences in size and shape of pollen and stigmas papillae (Dulberger et al. 1975; Dulberger 1992; Barrett and Shore 2008). However, it seems not always a valid association. A small number of HetSI species, like *Turnera joelii* and *Turnera scabra* (Turneraceae),, were already reported with similar pollen shape and papillae structure regardless of their floral morphs (Safavian and Shore 2010). Moreover, LSI systems are not restricted to heteromorphic species as it was also found in homomorphic species like in *Ipomopsis aggregate* from the Polemoniaceae (Sage et al. 2006). And when found in heteromorphic species, LSI systems can be disconnected from the segregation of the floral morphs, and thus classify as nonheteromorphic LSI systems. This is the case of three style-polymorphic Amaryllidaceae species, *Narcissus tazetta* L., *Narcissus triandrus* L. and *Narcissus papyraceus*. They reproduce controlled by LSI systems and present similar pollen shape and papillae structure between floral morphs. However, their LSI systems are not related to the heteromorphy of their flowers, as crosses between individuals of the same floral type are fertile, and only strict self-pollination results into ovarian prezygotic pollen rejection (Dulberger 1964; Sage et al. 1999; Barrett and Shore 2008; Simon-Porçar et al. 2015). For *Lgh*, we didn’t assess the genetic identity of the sampled individuals in our study along the 530km west-east transect of the Loire watershed. We were thus not able to identify if the LSI we identified was truly heteromorphic, or as, in *Narcissus sp*., nonheteromorphic.

### Occurrence of the prezygotic LSI system in the Myrtales order

In the order of Myrtales, 76 of 674 species of the Onagraceae family were reported to be self-incompatible (Raven 1979). Two of these species, *Oenothera organensis* and *Oenothera rhombipetala*, were formally demonstrated to be self-incompatible by studying their pollen tube progression after self and cross pollinations and were reported to involve a homomorphic gametophytic SI system (Emerson 1939; Bali and Hecht 1965). In *Epilobium obcordatum*, another species of the Onagraceae family suspected to mate using an ovarian, LSI system (Seavey and Bawa 1986), 38.6% of the self-pollen tubes stopped before entering the embryo sac (Seavey and Carter 1996). However, contrary to our observations in *Lgh*, the rare self-fertilised ovules of *E. obcordatum* present substantial rates of post-zygotic failures resulting in an average of ~5% of viable seed set (Seavey and Carter 1994).

Overall, LSI have been previously reported in two families of the Myrtales order: in the Myrtaceae family, in *Acca sellowiana* (Finatto et al. 2011), in *Melaleuca alternifolia* (Baskorowati et al. 2010) and in *Eucalyptus globulus* (Pound et al. 2002), and in six *Vochysia* species of the Vochysiaceae family (Oliveira and Gibbs 1994). If confirmed, our results may add a third family developing an LSI system in the Myrtales order, call for a potential reappraisal of the type of SI developed by *Oenothera organensis* and *O. rhombipetala* of the Onagraceae family as evidenced by Emerson (1939) and Ball &Hecht (1965) and question a possible wider occurrence of LSI in this order as reported in Gibbs (2014).

### Permeability of LSI L-morph flowers: a delayed selfed mating system?

Our results showed that, despite the LSI developed in the L-morph flowers, self-fertilisation still concerned 0.01% to 0.1% of the available ovules, in both experimental and *in natura* populations. This type of breeding system, named delayed selfing, enables self-fertilisation especially at the end of flowering season. It has been reported in multiple angiosperm species as a recurrent breeding strategy (Lloyd 1992; Sakai 1995; Goodwillie and Weber 2018). In such species, self-pollination is blocked or delayed until the opportunity for outcrossing has passed. Delayed selfing is thought to have evolved in allogamous species because it would provide reproductive assurance when populations suffer from the lack of compatible pollen (Goodwillie and Weber 2018; Ruane et al. 2020; Xu 2021).

### A preferential allogamy breeding system even in the self-compatible S-morph?

Before our study, we expected that individuals of the self-compatible S-morph preferentially mates using selfing (Kerbs et al. 2020). Surprisingly, the elongation dynamics of pollen tubes highlighted another process susceptible to favour allogamy in *Lgh*, even in the self-compatible S-morph. In both floral morphs, inter-morph pollen tubes elongated significantly faster than self-pollen tubes while progressing along the style. This phenomenon has already been described in several species with a LSI mating system, like in *Camellia oleifera* and in *Crotalaria juncea* (Liao et al. 2014; Rangappa Thimmaiah et al. 2018). But faster elongation of inter-morph pollen tubes does not occur in all LSI species. For example, self- and cross-pollen tubes present similar elongation speed in *Aconitum kusnezofii* or *Cyrtanthus breviflorus* (Vaughton et al. 2010; Hao et al. 2012). Further studies should determine whether the advantage of inter-morph pollen tube elongation also applies when flowers are pollinated with a mixture of self and inter-morph pollens in controlled pollinations, as reported in *Campsis radicans* (Bertin et al. 1989) or using paternity analysis on seeds produced after free pollination in *in natura* populations that mix the two floral morphs, as reported in *Luculia pinceana* (Zhou et al. 2015).

### Consequences of LSI for invasive populations of Lgh

Our results showed that invasive populations only composed of the self-incompatible L-morph still produced few small fruits containing few seeds resulting from self-fertilization, in similar proportions to what we obtained in experimental conditions. If these self-fertilized seeds remained limited in number at the scale of an individual, we estimated from our measures in quadrats that populations in Loire watershed still produced from 92 to 217 seeds per m^2^ which can all potentially germinate and resulted into viable seedlings. Delayed selfing in the L-morph *Lgh* populations resulted in a consequential quantity of viable floating seeds that may contribute to their local regeneration, seed bank and dissemination, provided that inbreeding depression would not affect such selfed regeneration at later stages. In the invasion context, individuals capable of self-fertilization or uniparental reproduction can settle a new population in the absence of compatible partners (Baker 1955; Barrett et al. 2008). Accordingly, a recent survey on 1,752 angiosperm species showed that selfing ability fosters directly and indirectly alien plant establishment (Razanajatovo et al. 2016). In contrast, 75% of the worldwide *Lgh* invasive populations are monomorphic L-morph self-incompatible populations (Portillo-Lemus et al. 2021a) which may constitute an anomaly to ‘Baker’s Law’ (Baker 1955). Our results showed that L-morph individuals certainly mated using a LSI system but also used delayed selfing in European invasive populations, providing support for Baker’s hypothesis. This breeding system with preferential allogamy and delayed selfing, never described before this study, must be considered for the future management plan of *Lgh* invasive populations.

Beyond a system with a compatible S-morph and a self-incompatible L-morph, that may have suggested simple breeding systems, our study revealed a more complex picture, where the two floral morphs combined different mechanisms that may result in preferential allogamy and delayed selfing. Our results encouraged to explore in more details fertilization in plants to better characterise their breeding systems and their consequences for populations. Concerning *Lgh*, we still lack of a clear picture of the respective importance of its reproductive modes in native and worldwide invasive populations in different ecological contexts. Especially, how its reproductive modes may explain and structure these monomorphic populations found along the recent European invasion front. Genetic marker-based analyses of experimental crosses and parentage analyses in natural populations should help validating the relevance of our first results and clarifying the reproductive modes of both floral morphs and natural populations as recently achieved in two Oleaceae species (Besnard et al. 2020, De Cauwer et al. 2021).

## Data accessibility

Data are available online at https://doi.org/10.5281/zenodo.5106805 (Portillo-Lemus et al. 2021b).

## Author Contribution

LP and DB designed this project. MH, MB, LP and DB performed all experiments. JH participated in capsule husking. LP, SS and DB analysed data and wrote the manuscript. SS and DB answered and made corrections required by the reviewing process. All authors approved the manuscript.

## Acknowledgements

We warmly thank Noni Franklin-Tong, Emiliano Mora-Carrera, Antoine Vernay, Juan Arroyo and two anonymous reviewers for their useful comments that helped shaping this manuscript. This research was supported by FEDER funds from Région Centre-Val de Loire and by Agence de l’eau Loire-Bretagne (grant Nature 2045, programme 9025 (AP 2015 9025)). FEDER also financed the doctoral grant of L. Portillo and the technical assistance salary of M. Harang. The authors thank Diane Corbin (FRAPNA Loire - Ecopôle du Forez), and Guillaume Le Roux (Réserve Naturelle Val d’Allier Châtel-de-Neuvre) for making plant material available. We thank the Experimental Unit of Aquatic Ecology and Ecotoxicology (U3E) 1036, Institut national de recherche pour l’agriculture, l’alimentation et l’environnement (INRAE, which is part of the research infrastructure Analysis and Experimentations on Ecosystems-France, for help with the maintenance of plants. Version 4 of this preprint has been peer-reviewed and recommended by Peer Community In Ecology (https://doi.org/10.24072/pci.ecology.100095).

## Supplementary materials

**Table S1:** Fruit-set, seed-set and germination rate in five S-morph and five L-morph monomorphic *in natura* populations in western Europe. Mean number of seeds per fruit (N_seeds/fruit_), fruit-set per square meter, seed-set per square meter produced in the studied *in natura* monomorphic populations of *Lgh*. Seed-set productions per m^2^ in monomorphic S-morph populations were estimated by counting the number of seeds produced by fruits in 5 quadrats of 1m^2^ and multiplied by the mean number of counted fruits per quadrat. Seed-set productions per 1m2 in L-morph populations are reported as the exhaustive number of seeds within all the fruits found in the quadrats of one square meter.

| Population | GPS Location | Floral morph | N<sub>seeds/fruit</sub> | Fruit-set/m<sup>2</sup> | Seed-set/m<sup>2</sup> | Germination (percentage) |
| --- | --- | --- | --- | --- | --- | --- |
| Chambéon | 45°41'03.4"N 4°12'12.1"E | L | 4.2 | 22 | 92 | 95.67 |
| Châtel-de-Neuvre | 46°24'05.4"N 3°19'10.1"E | L | 6.4 | 15 | 96 | 93.75 |
| Gilly-sur-Loire | 46°31'40.6"N 3°48'25.7"E | L | 6.8 | 28 | 190 | 93.68 |
| Pouilly-sur-Loire | 47°16'48.4"N 2°57'25.2"E | L | 5.7 | 19 | 108 | 89.81 |
| Orléans | 47°53'41.3"N 1°55'48.6"E | L | 6.2 | 35 | 217 | 94.47 |
| Pont-de-Cé | 47°25'40.7"N 0°31'28.9"W | S | 55.0 | 792 | 43560 | 90.30 |
| Mazerolles | 47°23'17.3"N 1°28'07.4"W | S | 60.0 | 864 | 51840 | 96.71 |
| Saint-Aignan-sur-Cher | 47°16'19.8"N 1°22'32.1"E | S | 55.0 | 528 | 29040 | 89.46 |
| Lac-de-Maine | 47°27'34.7"N 0°35'30.8"W | S | 50.0 | 624 | 31200 | 98.12 |
| Sabot-d'Or | 47°19'14.1"N 2°15'24.4"W | S | 50.0 | 384 | 19200 | 91.55 |

**Figure S1:**
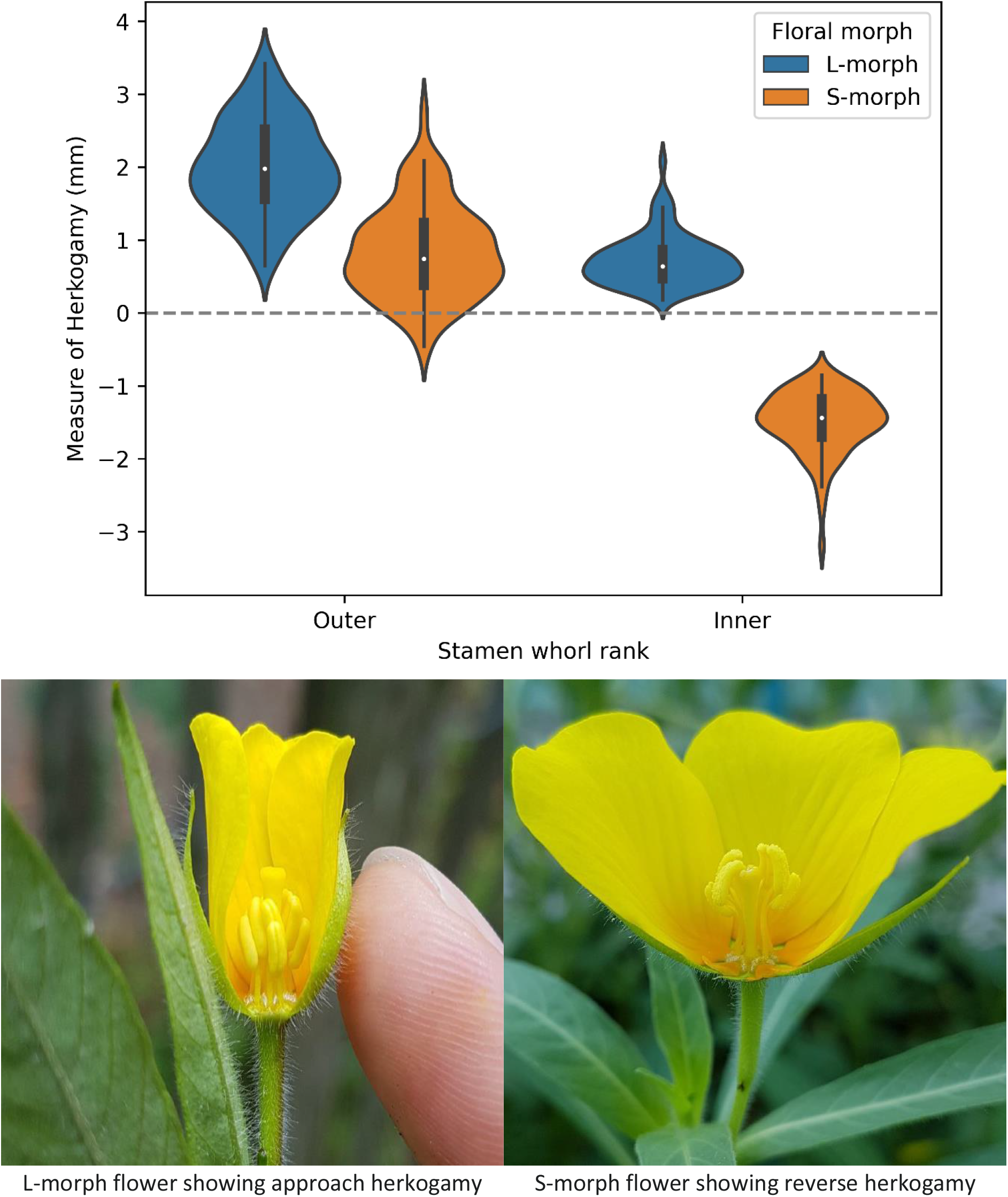
Measures and photos of herkogamy in self-incompatible L-morph and self-compatible S-morph flowers found in Western European populations of Lgh. Measures are reported for the outer and inner whorls of anthers. 150 measured flowers per violin plot, 5 populations sampled across the Loire watershed per morph.

**Figure S2:**
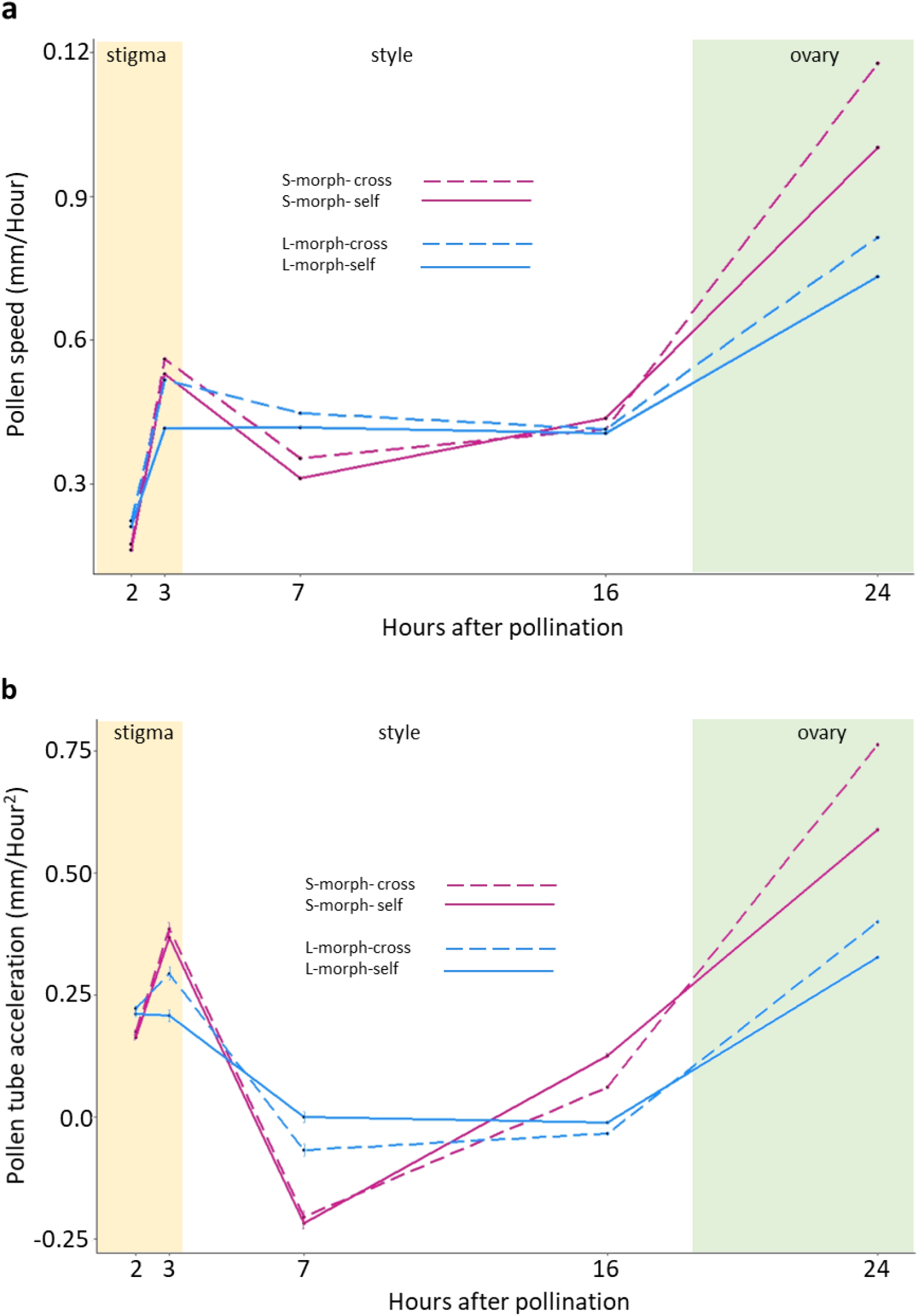
Speed and acceleration of pollen tube elongations in both floral morphs in *Ludwigia grandiflora* subsp. *hexapetala* after self- and cross-pollinated. Pink = S-morph, blue = L-morph; solid line = self-pollination (self) and dotted line=cross-pollination (cross). a) Speed of pollen tube elongations in both S-morph and L-morph styles after self and cross pollinations. b) Acceleration of pollen tube elongations in both S-morph and L-morph styles after self and cross pollinations.

**Table S2:**
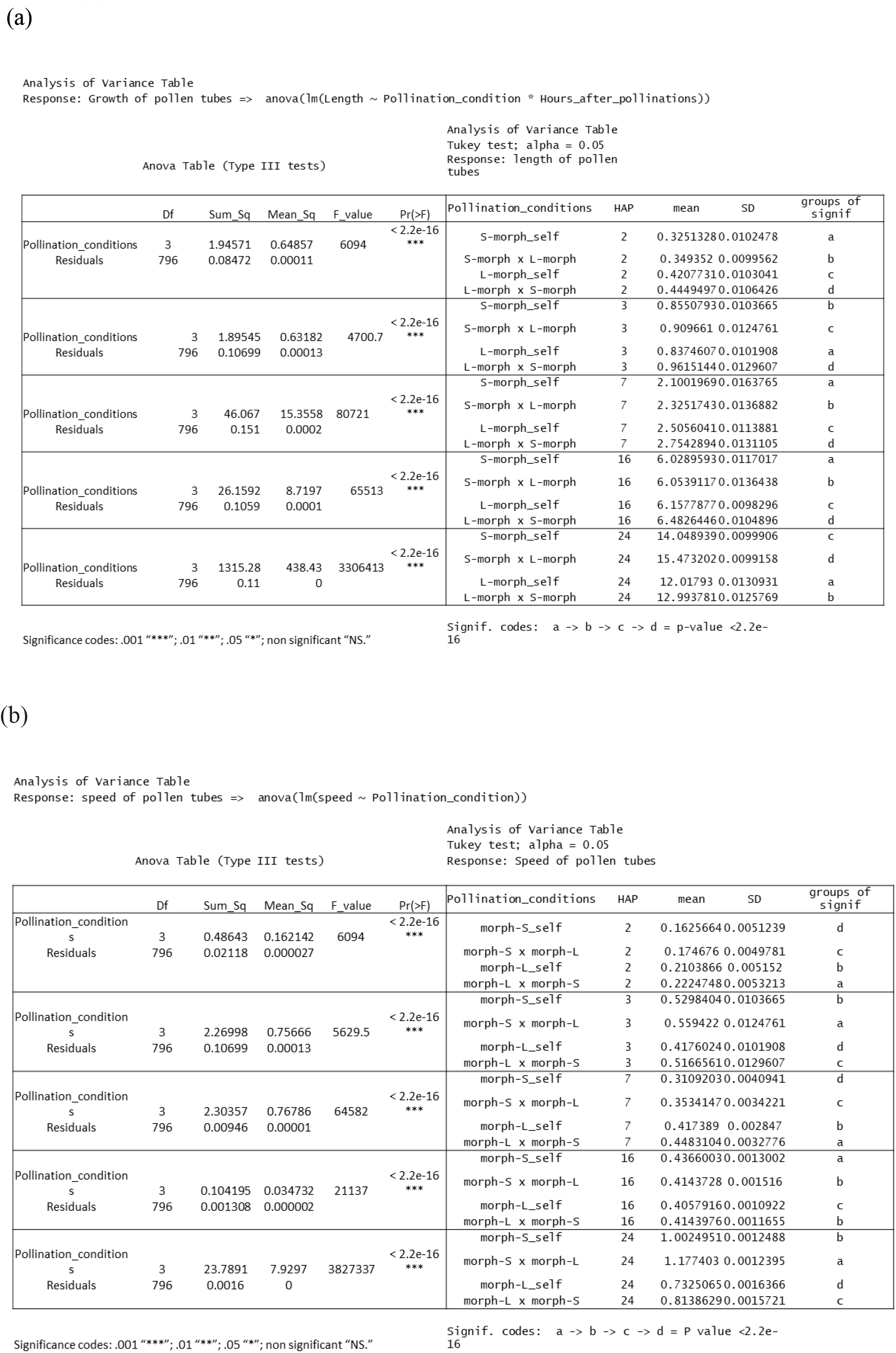
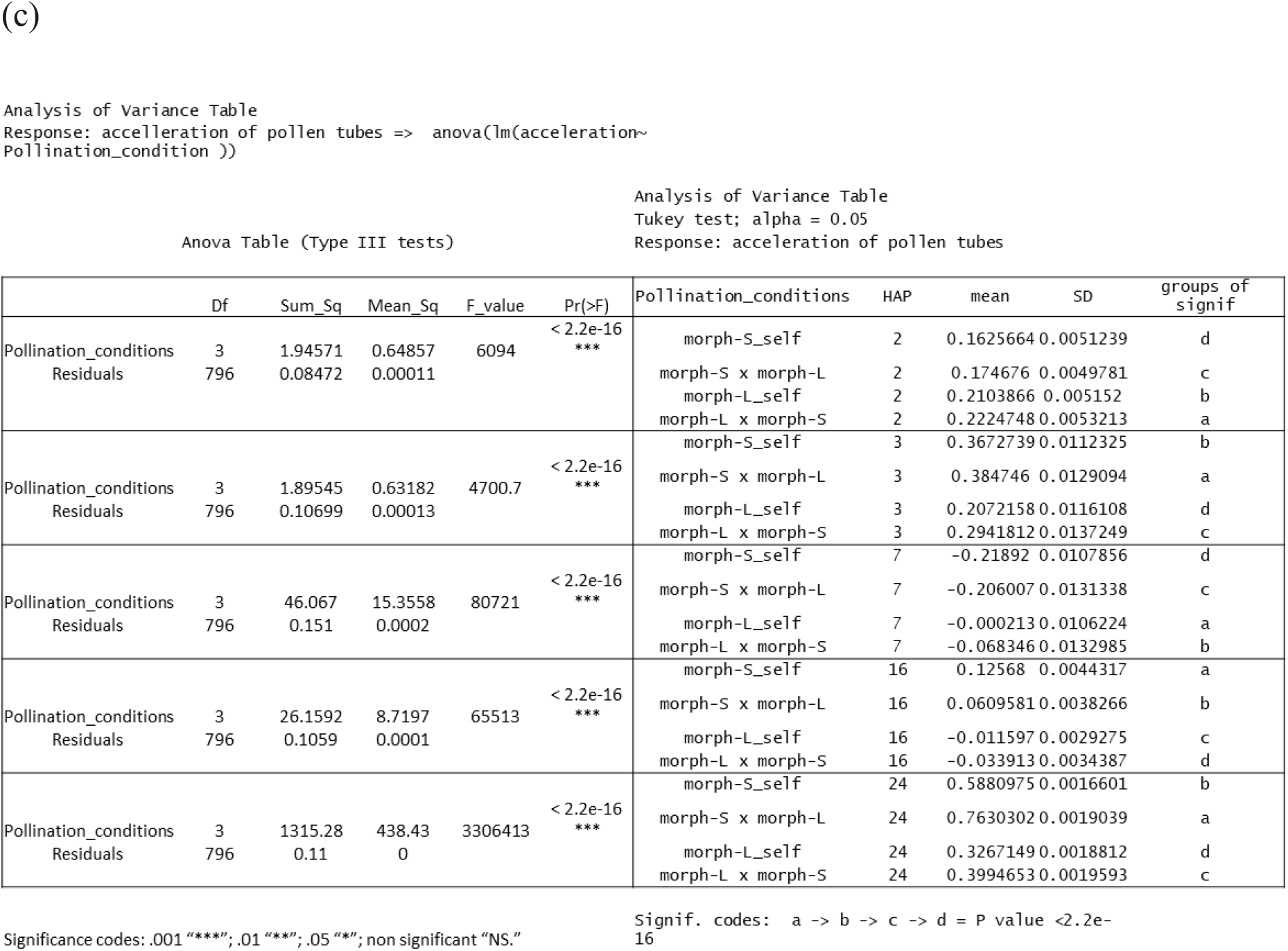
Analyses of the variations of pollen tube elongation along time (a), speed (b) and acceleration of pollen tube elongation (b) in both floral morphs in *Ludwigia grandiflora* subsp*. hexapetala* after self- and cross-pollinations.

|  | Df | Sum_Sq | Mean_Sq | F_value | Pr(>F) | Pollination_conditions | HAP | mean | SD | groups of signif |
| --- | --- | --- | --- | --- | --- | --- | --- | --- | --- | --- |
| Pollination_conditions | 3 | 1.94571 | 0.64857 | 6094 | < 2.2e-16 *** | S-morph_self | 2 | 0.32513280 | 0.0102478 | a |
| Residuals | 796 | 0.08472 | 0.00011 |  |  | S-morph x L-morph | 2 | 0.349352 | 0.0099562 | b |
|  |  |  |  |  |  | L-morph_self | 2 | 0.42077310 | 0.0103041 | c |
|  |  |  |  |  |  | L-morph x S-morph | 2 | 0.44494970 | 0.0106426 | d |
| Pollination_conditions | 3 | 1.89545 | 0.63182 | 4700.7 | < 2.2e-16 *** | S-morph_self | 3 | 0.85507930 | 0.0103665 | b |
| Residuals | 796 | 0.10699 | 0.00013 |  |  | S-morph x L-morph | 3 | 0.909661 | 0.0124761 | c |
|  |  |  |  |  |  | L-morph_self | 3 | 0.83746070 | 0.0101908 | a |
|  |  |  |  |  |  | L-morph x S-morph | 3 | 0.96151440 | 0.0129607 | d |
| Pollination_conditions | 3 | 46.067 | 15.3558 | 80721 | < 2.2e-16 *** | S-morph_self | 7 | 2.10019690 | 0.0163765 | a |
| Residuals | 796 | 0.151 | 0.0002 |  |  | S-morph x L-morph | 7 | 2.32517430 | 0.0136882 | b |
|  |  |  |  |  |  | L-morph_self | 7 | 2.50560410 | 0.0113881 | c |
|  |  |  |  |  |  | L-morph x S-morph | 7 | 2.75428940 | 0.0131105 | d |
| Pollination_conditions | 3 | 26.1592 | 8.7197 | 65513 | < 2.2e-16 *** | S-morph_self | 16 | 6.02895930 | 0.0117017 | a |
| Residuals | 796 | 0.1059 | 0.0001 |  |  | S-morph x L-morph | 16 | 6.05391170 | 0.0136438 | b |
|  |  |  |  |  |  | L-morph_self | 16 | 6.15778770 | 0.0098296 | c |
|  |  |  |  |  |  | L-morph x S-morph | 16 | 6.48264460 | 0.0104896 | d |
| Pollination_conditions | 3 | 1315.28 | 438.43 | 3306413 | < 2.2e-16 *** | S-morph_self | 24 | 14.0489390 | 0.0099906 | c |
| Residuals | 796 | 0.11 | 0 |  |  | S-morph x L-morph | 24 | 15.4732020 | 0.0099158 | d |
|  |  |  |  |  |  | L-morph_self | 24 | 12.01793 | 0.0130931 | a |
|  |  |  |  |  |  | L-morph x S-morph | 24 | 12.9937810 | 0.0125769 | b |

|  | Df | Sum_Sq | Mean_Sq | F_value | Pr(>F) | Pollination_conditions | HAP | mean | SD | groups of signif |
| --- | --- | --- | --- | --- | --- | --- | --- | --- | --- | --- |
| Pollination_condition | 3 | 0.48643 | 0.162142 | 6094 | < 2.2e-16 *** | morph-S_self | 2 | 0.16256640 | 0.0051239 | d |
| Residuals | 796 | 0.02118 | 0.000027 |  |  | morph-S x morph-L | 2 | 0.174676 | 0.0049781 | c |
|  |  |  |  |  |  | morph-L_self | 2 | 0.2103866 | 0.005152 | b |
|  |  |  |  |  |  | morph-L x morph-S | 2 | 0.22247480 | 0.0053213 | a |
| Pollination_condition | 3 | 2.26998 | 0.75666 | 5629.5 | < 2.2e-16 *** | morph-S_self | 3 | 0.52984040 | 0.0103665 | b |
| Residuals | 796 | 0.10699 | 0.00013 |  |  | morph-S x morph-L | 3 | 0.559422 | 0.0124761 | a |
|  |  |  |  |  |  | morph-L_self | 3 | 0.41760240 | 0.0101908 | d |
|  |  |  |  |  |  | morph-L x morph-S | 3 | 0.51665610 | 0.0129607 | c |
| Pollination_condition | 3 | 2.30357 | 0.76786 | 64582 | < 2.2e-16 *** | morph-S_self | 7 | 0.31092030 | 0.0040941 | d |
| Residuals | 796 | 0.00946 | 0.00001 |  |  | morph-S x morph-L | 7 | 0.35341470 | 0.0034221 | c |
|  |  |  |  |  |  | morph-L_self | 7 | 0.417389 | 0.002847 | b |
|  |  |  |  |  |  | morph-L x morph-S | 7 | 0.44831040 | 0.0032776 | a |
| Pollination_condition | 3 | 0.104195 | 0.034732 | 21137 | < 2.2e-16 *** | morph-S_self | 16 | 0.43660030 | 0.0013002 | a |
| Residuals | 796 | 0.001308 | 0.000002 |  |  | morph-S x morph-L | 16 | 0.4143728 | 0.001516 | b |
|  |  |  |  |  |  | morph-L_self | 16 | 0.40579160 | 0.0010922 | c |
|  |  |  |  |  |  | morph-L x morph-S | 16 | 0.41439760 | 0.0011655 | b |
| Pollination_condition | 3 | 23.7891 | 7.9297 | 3827337 | < 2.2e-16 *** | morph-S_self | 24 | 1.00249510 | 0.0012488 | b |
| Residuals | 796 | 0.0016 | 0 |  |  | morph-S x morph-L | 24 | 1.177403 | 0.0012395 | a |
|  |  |  |  |  |  | morph-L_self | 24 | 0.73250650 | 0.0016366 | d |
|  |  |  |  |  |  | morph-L x morph-S | 24 | 0.81386290 | 0.0015721 | c |
| Pollination_conditions | 3 | 1.94571 | 0.64857 | 6094 | < 2.2e-16 *** | morph-S_self | 2 | 0.1625664 | 0.0051239 | d |
| Residuals | 796 | 0.08472 | 0.00011 |  |  | morph-S x morph-L | 2 | 0.174676 | 0.0049781 | c |
|  |  |  |  |  |  | morph-L_self | 2 | 0.2103866 | 0.005152 | b |
|  |  |  |  |  |  | morph-L x morph-S | 2 | 0.2224748 | 0.0053213 | a |
|  |  |  |  |  |  | morph-S_self | 3 | 0.3672739 | 0.0112325 | b |
| Pollination_conditions | 3 | 1.89545 | 0.63182 | 4700.7 | < 2.2e-16 *** | morph-S x morph-L | 3 | 0.384746 | 0.0129094 | a |
| Residuals | 796 | 0.10699 | 0.00013 |  |  | morph-L_self | 3 | 0.2072158 | 0.0116108 | d |
|  |  |  |  |  |  | morph-L x morph-S | 3 | 0.2941812 | 0.0137249 | c |
|  |  |  |  |  |  | morph-S_self | 7 | -0.21892 | 0.0107856 | d |
| Pollination_conditions | 3 | 46.067 | 15.3558 | 80721 | < 2.2e-16 *** | morph-S x morph-L | 7 | -0.206007 | 0.0131338 | c |
| Residuals | 796 | 0.151 | 0.0002 |  |  | morph-L_self | 7 | -0.000213 | 0.0106224 | a |
|  |  |  |  |  |  | morph-L x morph-S | 7 | -0.068346 | 0.0132985 | b |
|  |  |  |  |  |  | morph-S_self | 16 | 0.12568 | 0.0044317 | a |
| Pollination_conditions | 3 | 26.1592 | 8.7197 | 65513 | < 2.2e-16 *** | morph-S x morph-L | 16 | 0.0609581 | 0.0038266 | b |
| Residuals | 796 | 0.1059 | 0.0001 |  |  | morph-L_self | 16 | -0.011597 | 0.0029275 | c |
|  |  |  |  |  |  | morph-L x morph-S | 16 | -0.033913 | 0.0034387 | d |
|  |  |  |  |  |  | morph-S_self | 24 | 0.5880975 | 0.0016601 | b |
| Pollination_conditions | 3 | 1315.28 | 438.43 | 3306413 | < 2.2e-16 *** | morph-S x morph-L | 24 | 0.7630302 | 0.0019039 | a |
| Residuals | 796 | 0.11 | 0 |  |  | morph-L_self | 24 | 0.3267149 | 0.0018812 | d |
|  |  |  |  |  |  | morph-L x morph-S | 24 | 0.3994653 | 0.0019593 | c |

Significance codes: .001 "\*\*\*\*"; .01 "\*\*\*"; .05 "\*\*"; non significant "NS."
Signif. codes: a -> b -> c -> d = P value < 2.2e-16

**Figure S3:**
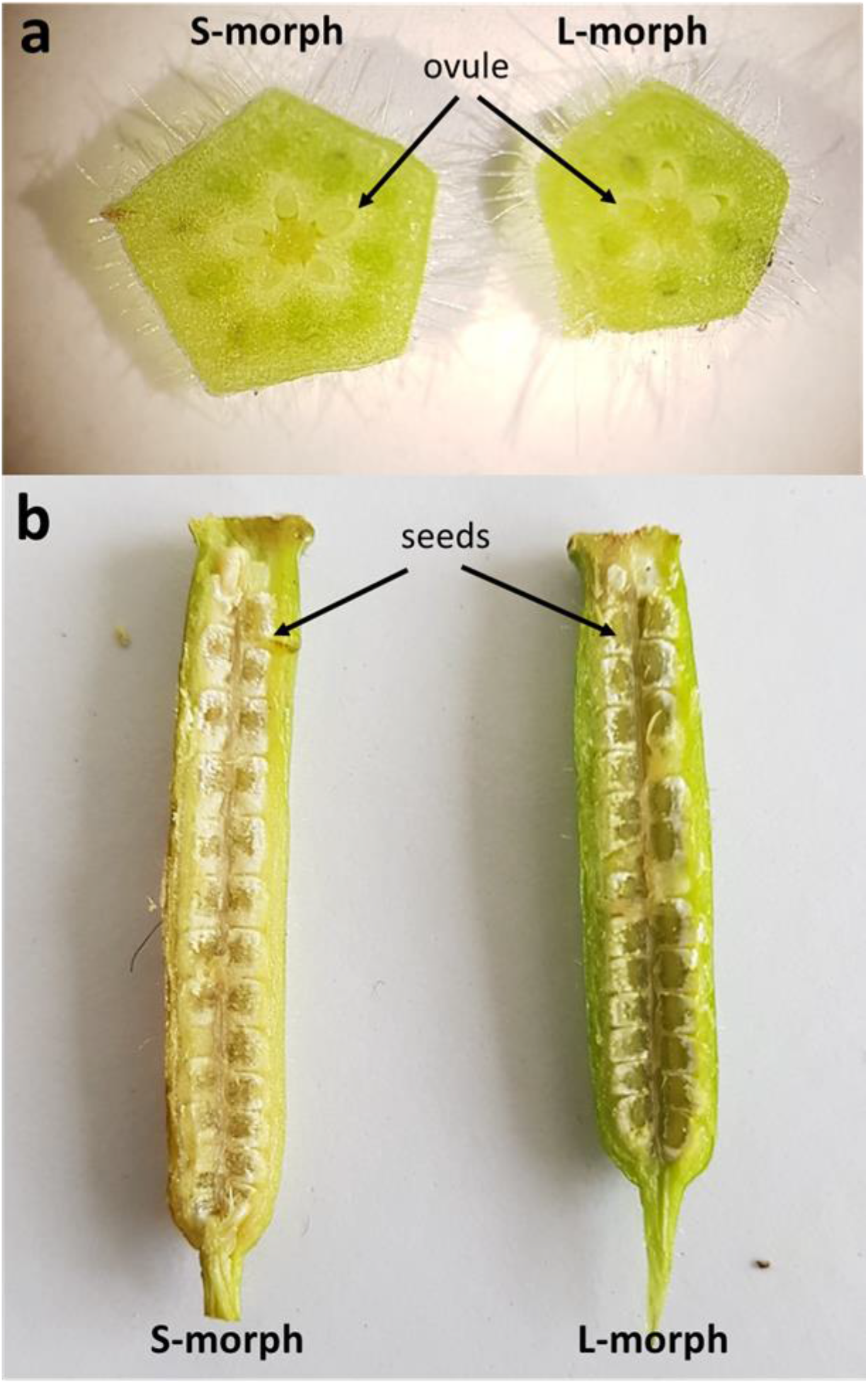
a) transversal and b) longitudinal sections of petaloculate ovary from 5-merous floral morphs of *Ludwigia grandiflora* subsp. *hexapetala*.

